# Progressive loss of independence in neuronal representations predicts cognitive decline

**DOI:** 10.64898/2026.06.28.734833

**Authors:** Drew E.G. Sheets, Douglas A. Ruff, Ramanujan Srinath, Keon S. Allen, John H. Morrison, Marlene R. Cohen

**Affiliations:** Department of Neurobiology and Neuroscience Institute, University of Chicago, IL, USA; California National Primate Research Center, University of California Davis, Davis, California, USA; Department of Neurology, School of Medicine, University of California Davis, Sacramento, California, USA

## Abstract

Intelligent behavior depends on the brain’s ability to represent multiple features of the environment simultaneously while keeping those representations independent^1,2^. Patients with Alzheimer’s disease often mix up objects, people, and events^3–7^, raising the possibility that disease mixes up the way that information is represented in the brain. Here we show that the independence of visual representations progressively breaks down during early stages of disease progression in a rhesus macaque model of Alzheimer’s disease and related dementias^8–10^. In visual area V4, representations of different visual features become progressively less independent, such that the representation of one feature is increasingly influenced by the value of another. We term this loss of independence neuronal feature confusion. This neuronal change predicts a specific behavioral consequence: because feature representations become less independent, preferences associated with one visual feature increasingly influence visually guided choices associated with other, independent features. Using an analogous image-selection task, we found the same behavioral signature in people with mild cognitive impairment, distinguishing them from age-matched controls. These results identify a specific and measurable alteration in neuronal population representations that predicts a behavioral change observed across species. More broadly, these findings demonstrate that neuronal population representations can guide the development of sensitive, non-invasive behavioral methods for early detection of functional changes associated with Alzheimer’s disease.

## Introduction

Alzheimer’s disease progression is thought to involve years of molecular and cellular change before overt cognitive impairment emerges^11–14^. During this period, disease-related alterations in brain function must already influence how neuronal populations represent and use information. Neuronal population activity provides a critical link between pathology and behavior because theories relating neuronal population representations to behavior make it possible to interpret functional changes in mechanistic terms. Yet most approaches to Alzheimer’s disease and related dementia (ADRD) focus either on molecular pathology or overt cognitive impairment, rather than on how disease alters neuronal population representations that shape behavior. Molecular biomarkers and neuroimaging can reveal pathology but are often invasive or expensive^15–21^, while behavioral deficits detected by standard cognitive assessments typically emerge only after substantial disease progression^11,22–24^. As treatments that slow cognitive decline become increasingly available^11,23,25–28^, there is growing need for sensitive and scalable approaches that can detect early functional changes associated with disease^11,14,23,29–31^.

Behavioral measurements could help address this challenge if they capture changes in how neuronal population activity contributes to behavior before overt impairment emerges. Most cognitive assessments focus on domains such as memory and executive function, where deficits are often partially masked by compensatory strategies and typically become measurable only after substantial disease progression^32–35^. In contrast, visually guided behaviors depend on rapid transformations between sensory representations and action, limiting opportunities for overt compensation. The primate visual system is therefore particularly well suited for linking neuronal population activity to behavior because relationships between visual information, neuronal population representations, and behavior can be measured quantitatively at both the neuronal and behavioral levels^36–41^. In addition, brain areas involved in visual processing are affected early in ADRD progression^42,43^, raising the possibility that subtle alterations in visual representations may produce sensitive behavioral signatures of disease-related functional change.

A central requirement of visual representations is that multiple features of an image remain represented independently, such that information about one feature does not systematically distort the representation of another. For example, the representation of an object’s color should not depend on its texture. In previous work, we found that this independence can transiently break down under high cognitive load (e.g. task switching), producing a phenomenon we term feature confusion, in which information about one visual feature influences the neuronal population representation of another^1,2^. This alteration in neuronal population representations gives rise to predictable changes in behavior: visually guided choices become increasingly influenced by relationships between features that are normally represented independently^1,2^. For instance, if one feature consistently biases behavior, choices may also become increasingly influenced by other features whose representations are no longer distinct from that feature, despite remaining independent in the images themselves. These observations suggest that the independence of feature representations may be particularly vulnerable to disruption and raise the possibility that similar changes could emerge during early stages of neurodegenerative disease.

We therefore asked whether feature confusion emerges during early stages of ADRD. To test this, we used a rhesus macaque model of Alzheimer’s disease in which tauopathy is induced by bilateral injection of an adeno-associated virus expressing a double-mutant form of human tau into entorhinal cortex^8,9^. Pathology in this model spreads according to synaptic distance^8,9^ and produces early functional changes in the visual system^10^. Because neuronal population activity and behavior can be measured longitudinally in the same individuals throughout disease progression, this model provides experimental access to early functional changes that are difficult to study directly in humans. We recorded neuronal population activity in visual area V4 while monkeys performed an image-selection task and quantified the extent to which representations of different visual features remained independent over time.

Representations of visual features in V4 became progressively less independent during disease progression, despite preserved tuning for visual features. This change in neuronal population representations predicted systematic alterations in visually guided behavior, such that preferences associated with one feature increasingly influenced choices associated with other, independent features. Using an analogous image-selection task in humans, we identified the same behavioral signature in individuals with mild cognitive impairment. Together, these results identify a specific alteration in neuronal population representations that links disease progression to behavior across species and suggest that visually-guided behavior can reveal early functional changes associated with Alzheimer’s disease.

## Results

### Image feature representations in V4 remain intact but become increasingly non-independent during disease progression

To study how visual representations change during disease progression, we induced a recently established macaque model of Alzheimer’s disease by injecting an adeno-associated virus (AAV) expressing a double-mutant form of human tau into entorhinal cortex (ERC) of two rhesus monkeys (Macaca mulatta; adult males, 10 and 12 kg)^8,9^. Histological analyses confirmed accurate targeting of ERC, induction of tau pathology, and its spread over the course of the experiment^10^. Tau was rarely observed in neuronal cell bodies in V4 even after one year, suggesting that changes in V4 activity likely arise from altered corticocortical input activity rather than local degeneration. Biomarker measurements from plasma (pTau231/pTau181) independently confirmed progressive tau accumulation^10^.

At the time of virus injection, we implanted multielectrode arrays in visual area V4 to record neuronal population activity (several dozen neurons per session) throughout disease progression. When monkeys fixated centrally, the recorded neurons responded to images presented in the lower-right visual field that covered their joint receptive fields (Figure 1A, left). These neurons showed transient visually evoked increases in firing rate following image presentation (Figure 1A, right). Over approximately thirteen months, each monkey viewed approximately 3,000 natural images spanning a wide range of objects and scenes^44,45^ while performing the image-selection task described below.

**Figure 1.**
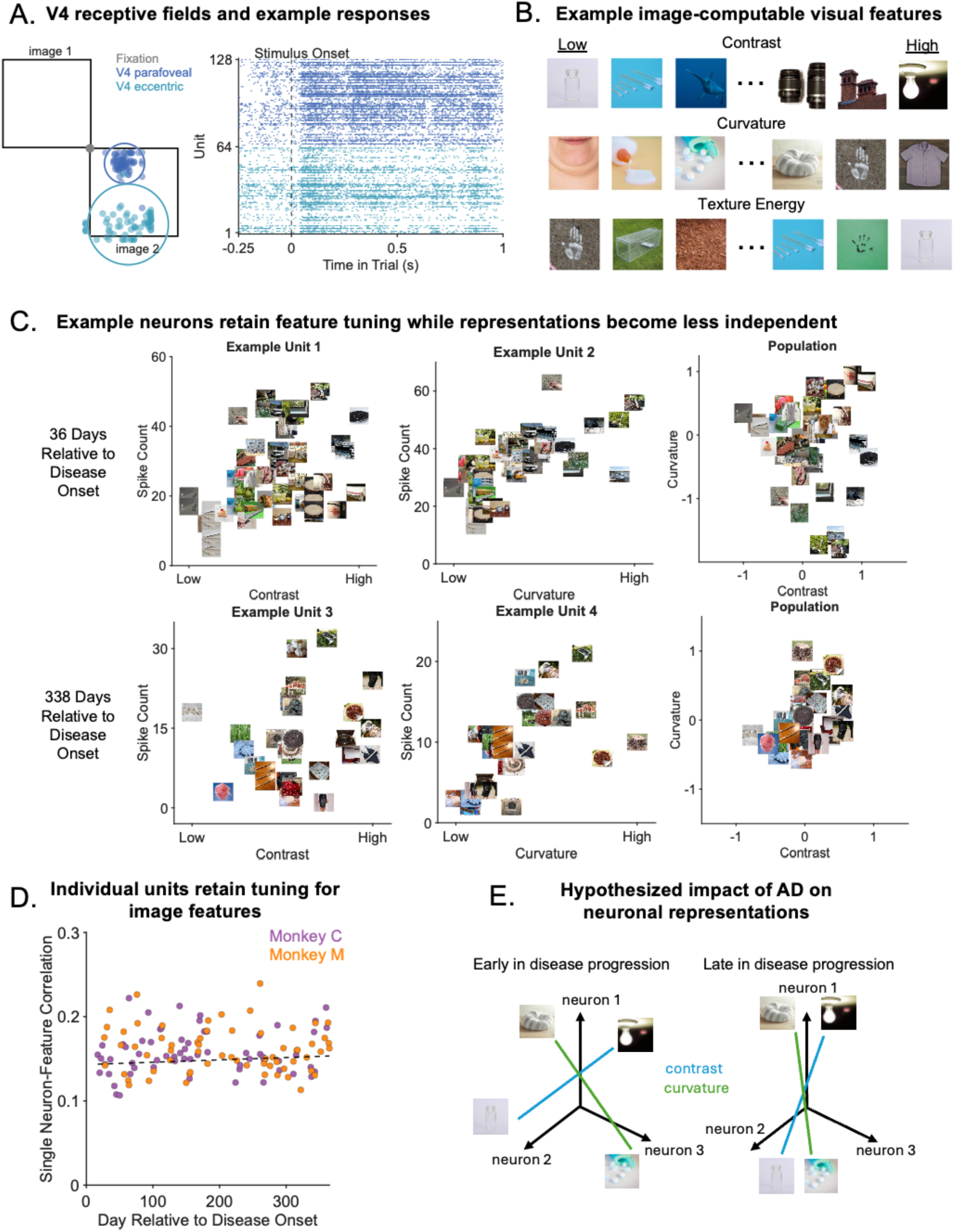
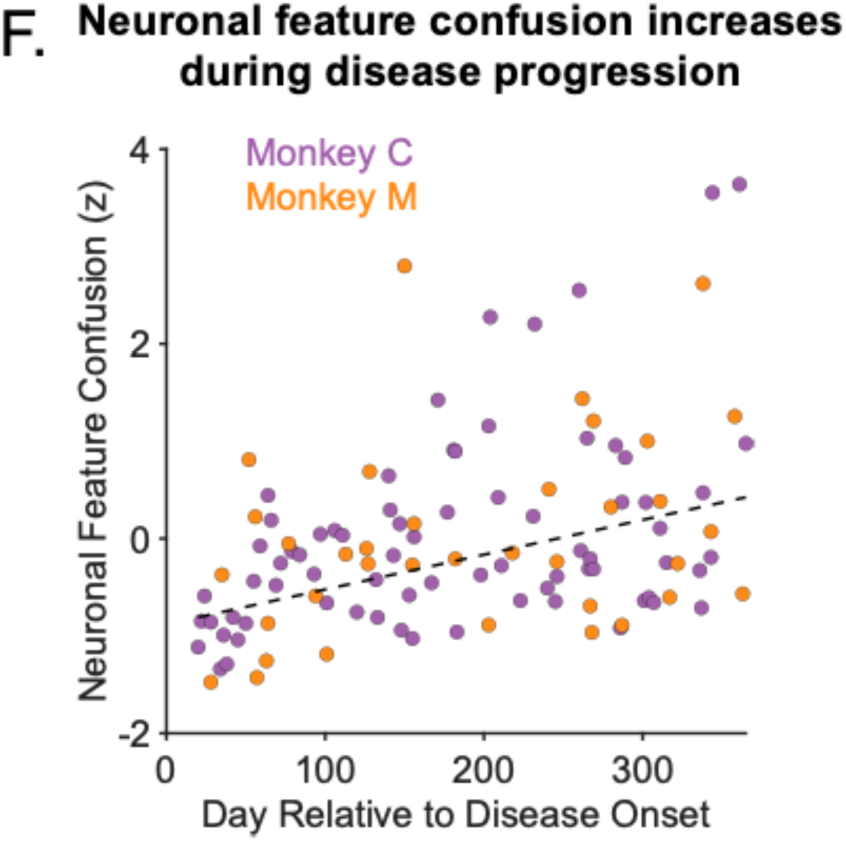
V4 representations of image features become non-independent during disease progression. (A) We chronically implanted two microelectrode arrays (Blackrock Microsystems, 64 electrodes per array) in mid-level visual cortex area V4 of each animal. The receptive fields (RF) of the recorded neurons covered parafoveal and more eccentric regions of the lower visual field. We recorded responses to pairs of images while monkeys fixated for at least 150 ms. One image was placed in the joint RFs of the recorded V4 units and the other was in the opposite hemifield. Neuronal analyses were based on responses during the 150 ms period of stable fixation. Each point denotes the center of the RF of an individual unit from the corresponding array. Each circle is the average RF center and median RF size for each array (blue, parafoveal array; tiel, eccentric array). (B) Example images spanning the range of three representative visual features: contrast, curvature, and texture. Images with the lowest (left) and highest (right) values for each feature are shown. (C) Individual V4 units remain selective for visual features throughout disease progression. Example neurons tuned to contrast (left) and curvature (middle) are shown early (day 36) and late (day 338) in disease progression. In contrast, population representations of these same features become less independent over time. For illustration, we identified the axes in neuronal population space that best represented contrast and curvature using QR decomposition. Early in disease progression, these axes were approximately orthogonal (day 36; n = 58 images, r = -0.11, p = 0.40), indicating independent feature representations. Later in disease progression, the axes became increasingly non-orthogonal (day 338; n = 29 images, r = 0.37, p = 0.048), indicating reduced independence between feature representations. (D) Tuning of individual units to image features remains stable over time. Image features were orthogonalized via principal components analysis and the variance bias was removed from each component via z-scoring. Correlations were computed between spike counts for each unit and values of a given feature dimension across all trials in a single session. Individual unit correlations were averaged across all features in a single session separately for each monkey. Throughout disease progression the ability of individual units to encode information about independent image features remained stable (n = 127 sessions across two monkeys, r = 0.032, p = 0.72). (E) Schematic illustrating neuronal feature confusion. Each schematic shows neuronal population representations of contrast (blue) and curvature (green) in a space where each dimension represents the response of one neuron. When feature representations are independent, the corresponding axes are orthogonal (left). Disease progression increases alignment between these axes (right), such that variation in one feature increasingly influences the representation of another. (F) Neuronal feature confusion increases during disease progression. Dimensionality reduction was applied to the image-computable features to orthogonalize feature dimensions and remove inherent correlations between image features. Principal components were z-scored to remove variance bias toward the first components. Cross-validated linear regression was used to identify the axis in V4 population space that best represented each feature dimension. Neuronal feature confusion (y-axis) was quantified as the ratio between cross-decoding performance (correlation between the actual values of one feature and projections onto the axis corresponding to another feature) and mean self-decoding performance for the feature pair. If feature representations are orthogonal, cross-decoding should approach zero. Neuronal feature confusion increased progressively throughout disease progression (the correlation between neuronal feature confusion on each session and time was significantly greater than 0; n = 110 sessions across two monkeys, r = 0.40, p = 1.6 x 10^-5^).

To relate neuronal population activity to properties of the images, we characterized each image using a set of image-computable visual features, including contrast, curvature, and texture energy (Figure 1B; Supplementary Figure 1; see Methods). These features allowed us to quantify how neuronal population responses depended on multiple features of the images. Because these features were extracted from natural images, the features were non-independent (Supplementary Figure 2) and often non-normally distributed across images. For this reason, for some analyses, we did dimensionality reduction on the image space (where each image is a point in the space and each dimension is the value of one feature; see Methods). This allowed us to investigate neuronal and behavioral correlates of features that are independent in the image space.

Throughout disease progression, individual V4 neurons continued to encode visual features. Example neurons remained selective for contrast and curvature both early and late in disease progression (see examples in Figure 1C, left and middle columns). Across the recorded neurons, tuning for image-computable features remained robust throughout the experiment (Figure 1D). Thus, disease progression did not eliminate feature information at the level of individual neurons.

We next asked whether the relationships among feature representations changed during disease progression. A central requirement of neuronal population representations is that independent visual features remain represented independently, such that variation in one feature does not systematically influence the representation of another. In previous work, we found that this independence can transiently break down under high cognitive load, producing feature confusion in which information about one feature influences representations of another^1,2^. We therefore asked whether a similar loss of independence between feature representations emerges during disease progression.

To test this, we analyzed neuronal population activity in a common population response space in which the response of each neuron defined one dimension. Within this space, we identified axes corresponding to individual visual features and examined whether neuronal population responses along those axes remained independent. Early in disease progression, projections of neuronal population activity onto axes corresponding to visual features that varied independently across images were approximately orthogonal, indicating that variation in one feature did not systematically influence the representation of another (for example, see Figure 1C, right column, top). Later in disease progression, these axes became increasingly aligned, indicating that representations of different features became less independent (Figure 1C, right column, bottom).

We define this loss of independence between feature representations as neuronal feature confusion (Figure 1E). When feature representations are independent, axes corresponding to different features remain orthogonal, such that a decoder trained on one feature should not generalize to another (Figure 1E, left). When feature representations become less independent, these axes become increasingly aligned, such that readout of one feature is systematically influenced by the value of another (Figure 1E, right).

We quantified neuronal feature confusion using cross-decoding of visual feature representations. After reducing the dimensionality of the image-computable features, we trained cross-validated linear decoders to predict each feature dimension from neuronal population activity in V4. We then tested whether a decoder trained to predict one feature dimension could also predict the value of a different feature dimension. If two features are represented independently, a decoder trained on one feature should not generalize to another. In contrast, if feature representations are not independent, cross-feature decoding should exceed chance levels. We quantified neuronal feature confusion for each pair of features as cross-decoding performance normalized by self-decoding performance, which provides an upper bound on decoding accuracy.

Together, these results indicate that disease progression alters the relationships among visual feature representations without eliminating feature information itself. Neuronal population representations of different visual features became progressively less independent during disease progression (Figure 1F). When combined with the observation that the encoding of individual features by individual neurons remained largely unchanged, these results suggest that the dominant change associated with disease progression was not loss of feature selectivity but a progressive increase in neuronal feature confusion at the population level.

### An image selection task as a probe for changes in visual cognition

We hypothesized that increased neuronal feature confusion in visual cortex during disease progression would alter visually guided behavior in a predictable way. If neuronal representations of distinct visual features become less independent, then the influence of one feature on behavior should increasingly depend on the value of another. For example, if a monkey preferentially selects images with high contrast, and neuronal representations of contrast and curvature become less independent, then curvature may increasingly influence behavior associated with contrast, even when the two feature dimensions are independent in the images themselves.

To test this prediction, we measured image selection behavior throughout disease progression using an image selection task adapted from established paradigms for studying novelty preference and visual memory^46–49^. Each trial began with central fixation, followed by presentation of two images on opposite sides of the screen. The monkeys maintained fixation for 150 ms while we measured neuronal responses (Figure 1A). After this brief exposure, the fixation cue disappeared and the monkeys freely viewed the images for 1000–1500 ms while eye position was monitored (Figure 2A). Monkeys were rewarded after the viewing period regardless of which image they selected. We defined the selected image on each trial as the image the monkey first fixated during the free-viewing period.

**Figure 2.**
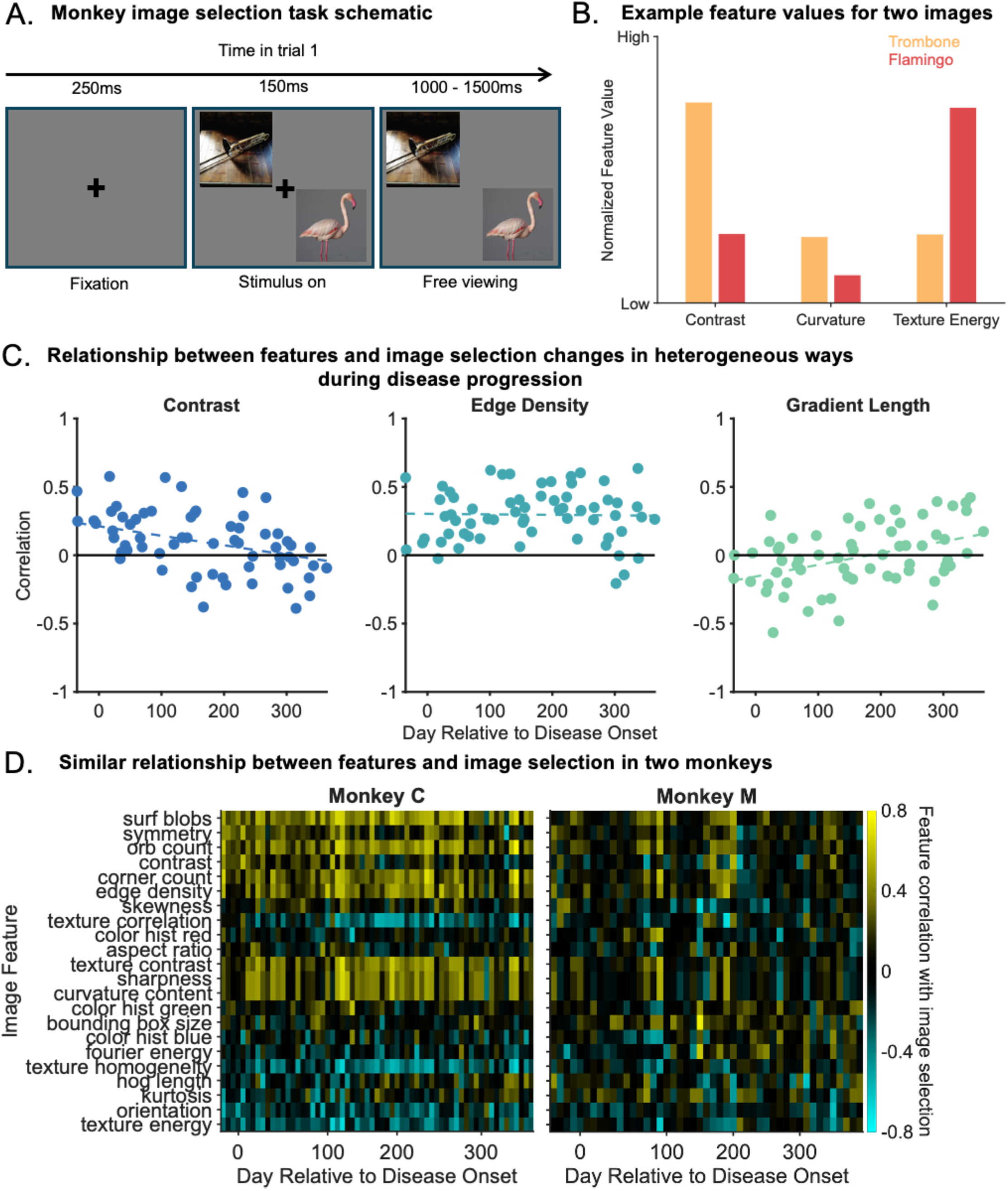
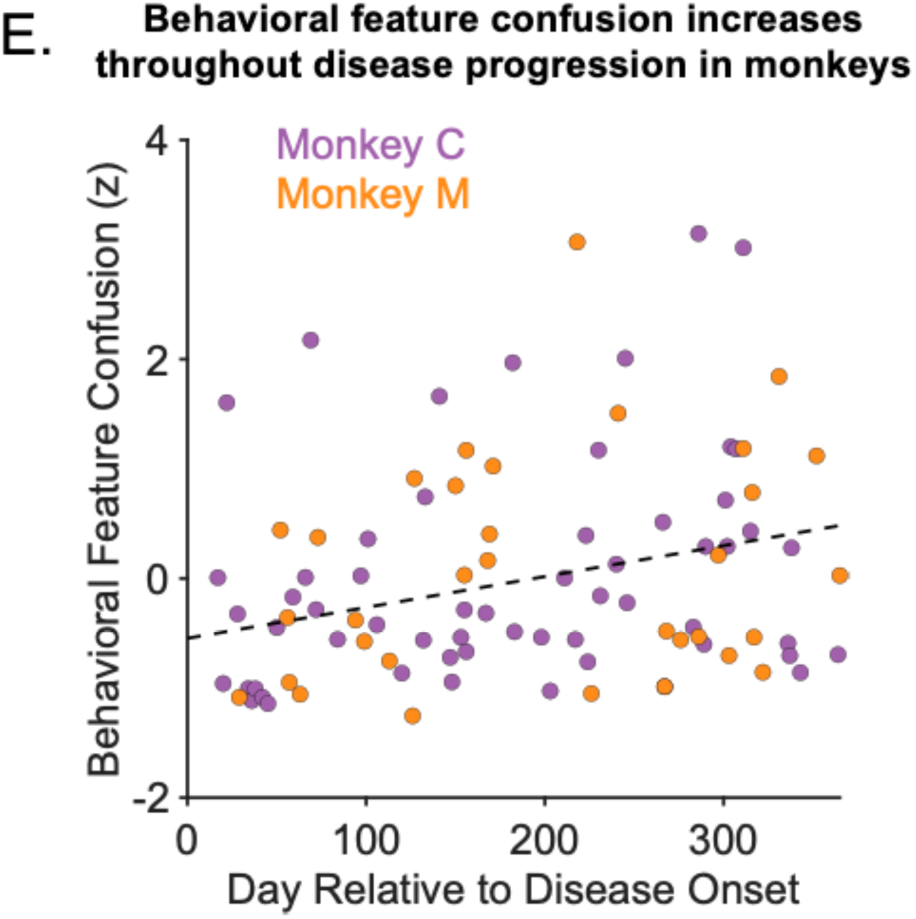
Behavioral feature confusion increases during disease progression. (A) Schematic of the monkey image selection task. Each trial began when the monkey fixated centrally for 250 ms, after which two images appeared on opposite sides of the screen. The lower-right image was positioned within the joint receptive fields of the recorded V4 neurons. After a 150 ms enforced fixation period during which neuronal responses were measured, the fixation point disappeared and the monkeys freely viewed either or both images for 1000–1500 ms to earn a reward. (B) Example image feature profiles for the two images shown in A. Differences in feature values between the selected and non-selected images were used to quantify relationships between image features and behavior. Positive relationships indicate that the image with the higher value of a feature was more likely to be selected, whereas negative relationships indicate that the image with the lower value of a feature was more likely to be selected. (C) Example relationships between image-computable features and image selection behavior over disease progression. Relationships between individual features and behavior could decrease (contrast; n = 63 sessions for Monkey C, r = -0.48, p = 7.3 x 10^-5^), remain stable (edge density; n = 63 sessions for Monkey C, r = 0.0054, p = 0.97), or increase (gradient length; n = 63 sessions for Monkey C, r = 0.40, p = 0.0011) over time. These heterogeneous changes indicate that disease progression did not simply reduce the overall influence of visual features on behavior. (D) Similar feature-based image selection behavior across monkeys. Both monkeys showed similar positive (yellow) and negative (cyan) relationships between image-computable features and behavior, including positive relationships for some features (e.g. orb count) and negative relationships for others (e.g. texture energy). (E) Behavioral feature confusion increases during disease progression. Dominant image-computable feature dimensions were extracted using principal components analysis and z-scored, producing feature dimensions that were independent in the images themselves. Each feature dimension was fit to image choice using logistic regression on a session-by-session basis. Behavioral feature confusion was quantified as the correlation between predictions for pairs of feature dimensions normalized by the average correlation between each prediction and actual image choice. This metric increased progressively throughout disease progression (slope on a session-by-session basis for both monkeys significantly different from 0; n = 99 sessions across two monkeys, r = 0.28, p = 0.0048), paralleling the increase in neuronal feature confusion shown in Figure 1E. Together, these results indicate that disease progression was associated with a change in the relationships between feature influences on behavior in parallel with changes in neuronal population representations.

We used image selection behavior to quantify how visual feature dimensions influenced choice. For each image pair, we compared feature values between the selected and non-selected images and measured the relationship between each feature dimension and image choice across trials within a session. For example, a trombone image may contain relatively high values along one feature dimension and low values along another, whereas a flamingo image may show the opposite pattern (Figure 2B). If the monkey consistently selects images with similar dominant features relative to the image they are paired with, this suggests that those feature dimensions influence image selection behavior. Across many trials, this approach allowed us to estimate the influence of each feature dimension on visually guided behavior.

Relationships between feature dimensions and image selection varied across sessions and across feature dimensions (Figure 2C). Some feature dimensions showed increasingly strong relationships with behavior over time, whereas others remained stable, indicating that disease progression did not simply reduce the overall influence of visual information on behavior. To compare feature–behavior relationships across animals and sessions, we visualized the correlation between each feature dimension and image selection on a session-by-session basis (Figure 2D). Both monkeys showed broadly similar patterns of feature-based image selection across disease progression, including stable preferences for higher values of some feature dimensions and lower values of others.

We next asked whether neuronal feature confusion predicted corresponding changes in visually guided behavior. Because these features were orthogonalized, any non-independence of their influence on behavior reflects mixing up of those features, rather than statistics of the images. We therefore hypothesized that neuronal feature confusion would produce a corresponding behavioral consequence: feature dimensions that became less independent in neuronal population representations would also have less independent influences on image selection behavior.

Paralleling the neuronal analysis, we quantified how strongly each feature dimension predicted choice within a session (self-prediction) and how strongly the predictions of one feature dimension generalized to another (cross-prediction). We defined behavioral feature confusion as cross-prediction normalized by self-prediction, averaged across feature pairs. Normalizing by self-prediction expresses confusion as a fraction of each pair’s predictive power, reflecting shared structure between feature dimensions rather than their raw strength.

Behavioral feature confusion progressively increased during disease progression (Figure 2E). Thus, the breakdown in the independence among neuronal feature representations was accompanied by a corresponding breakdown in the independence of feature influences on behavior. Together, these results show that neuronal feature confusion in visual cortex predicts systematic and measurable changes in visually guided behavior.

### Image selection patterns in humans reflect cognitive impairment

Our results indicate disease progression in monkeys produces changes in visually guided behavior that are predicted by neuronal feature confusion: influences of visual feature dimensions on behavior become progressively less independent. If similar behavioral signatures are present in humans, they could provide a non-invasive behavioral marker of early functional changes associated with cognitive impairment. We therefore asked whether humans with mild cognitive impairment (MCI) show the same reduction in the independence of feature influences on behavior observed during disease progression in monkeys. To test this, we adapted the image selection task for online testing in human participants (Figure 3A).

**Figure 3.**
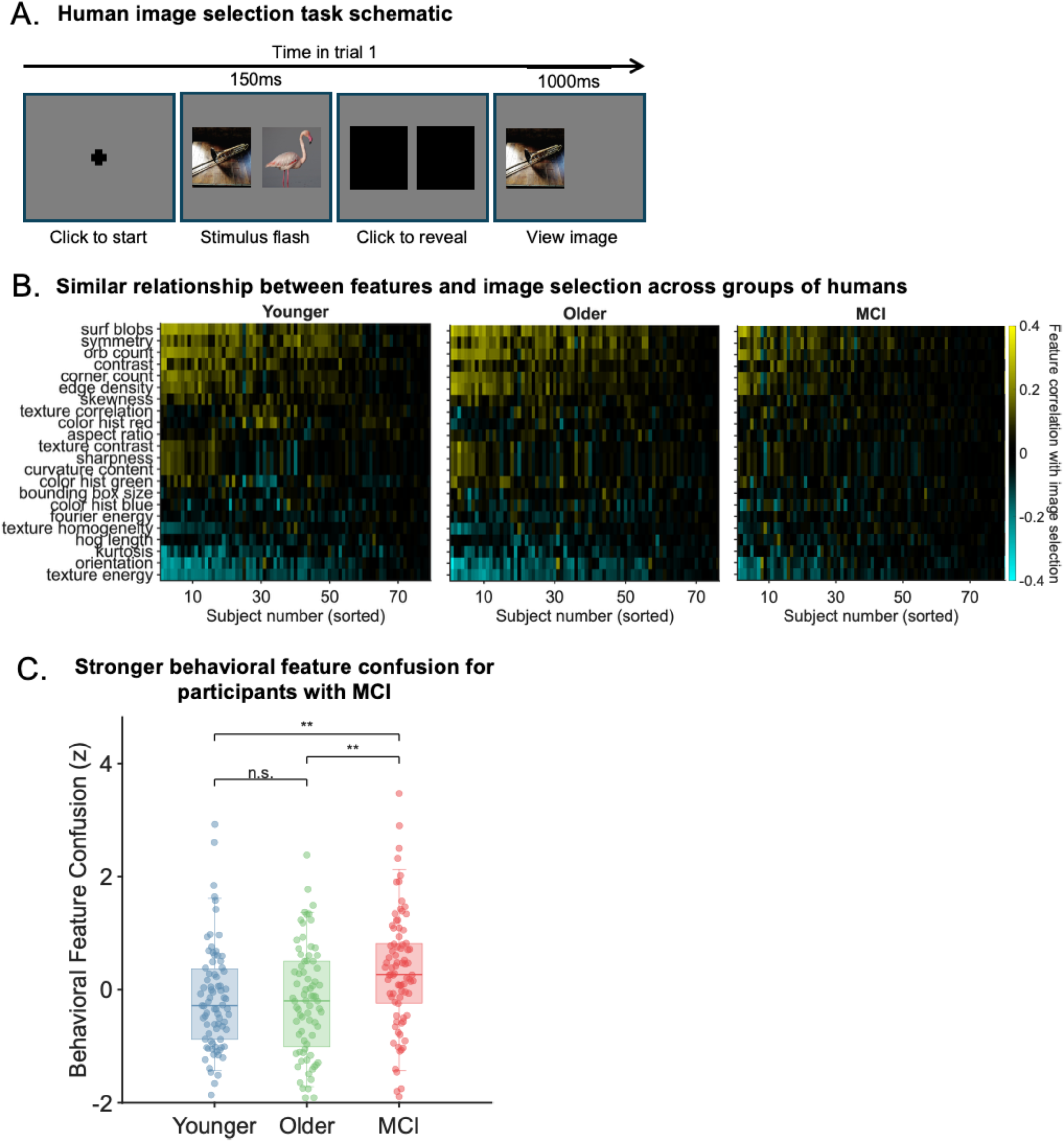
Human image selection behavior reveals behavioral signatures of feature confusion. (A) Schematic of the human online image selection task. Participants initiated each trial with a central mouse click, viewed two briefly presented images (150 ms), then selected one to reveal for longer viewing (1000 ms). (B) Relationships between visual features and image selection behavior in humans. For each participant, we compared the difference between the feature values in the left and right images on each trial and related those differences to image choice. Positive correlations (yellow) indicate preferential selection of the image with the higher feature value, whereas negative correlations (cyan) indicate preferential selection of the image with the lower feature value. Human participants showed feature-behavior relationships that were broadly similar to those observed in monkeys. Each column represents one participant, ordered according to the magnitude of the relationship between image features and choice (the same order as in the analogous analysis in monkeys; Figure 2D). (C) Behavioral feature confusion across human participant groups. Behavioral feature confusion did not differ between younger and older healthy adults (t-test, n = 79 younger vs. 76 older subjects, mean difference = 0.034, p = 0.82), but was significantly higher in individuals with MCI than in healthy controls (n = 155 healthy vs. 83 MCI subjects, mean difference = -0.50, p = 1.9 x 10^-4^). We used principal components analysis to reduce the dimensionality of the image features and z-scored the resulting components, producing feature dimensions that were independent in the images themselves. We fit logistic regression models to predict each subject’s image choices as a linear combination of feature dimensions and interactions. We quantified behavioral feature confusion as the ratio between the mean squared sum of interaction and linear weights. Boxes represent quartiles and stars represent significant differences (t-tests, significant differences were younger controls vs. MCI subjects, p = 0.0019; older controls vs. MCI, p = 0.0013).

Each trial began when the participant clicked a central fixation cross, after which two images were briefly presented side-by-side for 150 ms. The images then disappeared, and participants selected one image to reveal for longer viewing for 1000 ms. As in the monkey data, we quantified the correlation between the image-computable feature values and image selection behavior across trials for each participant.

Human participants showed feature-behavior relationships that were strikingly similar to those observed in monkeys (Figure 3B). Visual features that were positively associated with image selection in monkeys tended to be positively associated with image selection in humans, whereas features that were negatively associated with image selection in monkeys tended to show the same relationship in humans. Each column in Figure 3B represents one participant, ordered according to the magnitude of the relationship between image features and choice. Positive correlations (yellow) indicate preferential selection of images with higher values of a visual feature, whereas negative correlations (cyan) indicate preferential selection of images with lower values.

We next asked whether the independence of feature influences on behavior changed in individuals with MCI. As in the monkey analyses, we used principal components analysis to orthogonalize and reduce the dimensionality of the image-computable visual features, producing feature dimensions that were independent in the images themselves. We then predicted image choice by fitting logistic regression models with first-order terms and pairwise interactions between the orthogonalized feature dimensions. This allowed us to ask whether the influence of one feature dimension on behavior depended on the values of others. We quantified behavioral feature confusion in human subjects as the ratio of interaction term weights to first-order term weights (see Methods).

Behavioral feature confusion did not differ between younger and older healthy adults, but individuals with MCI showed significantly elevated behavioral feature confusion (Figure 3C). Thus, individuals with MCI exhibited the same fundamental behavioral signature predicted by neuronal feature confusion in the monkeys. Furthermore, healthy aging alone did not substantially alter the structure of feature influences on behavior, meaning that these measures hold promise for selectively identifying cognitive impairment.

Together, these results show that changes in the relationships among feature influences on behavior emerge in humans with mild cognitive impairment and parallel changes in neuronal population representations during disease progression in monkeys. These findings suggest that visually guided behavior may provide a sensitive behavioral readout of early functional changes associated with cognitive impairment.

### Neuronal population measurements can guide behavioral methods for detecting cognitive impairment

The results above show that image selection behavior differs in individuals with mild cognitive impairment and that these differences follow the same feature-based logic observed in monkeys later in disease progression. Humans respond to higher level image properties in ways that likely differ from monkeys. We therefore asked whether feature confusion in participants with MCI extends beyond image-computable visual features to features derived from human psychophysics experiments, including memorability, salience, and arousal. We characterized each image using a combination of low-level image features, psychophysics-derived measures (Supplemental Figure 3), and activations of a deep convolutional neural network (VGG-16^50^). Because these features are correlated in natural images, we used principal components analysis to identify orthogonal feature dimensions (Figure 4A). We then used the same method for computing behavioral feature confusion measure, fitting logistic regression models for each participant that predict image choice using linear combinations of feature dimensions and interactions. As before, we defined behavioral feature confusion in this richer feature space as the ratio of the sum of squared weights on the interaction and linear terms of the model (see Methods).

**Figure 4.**
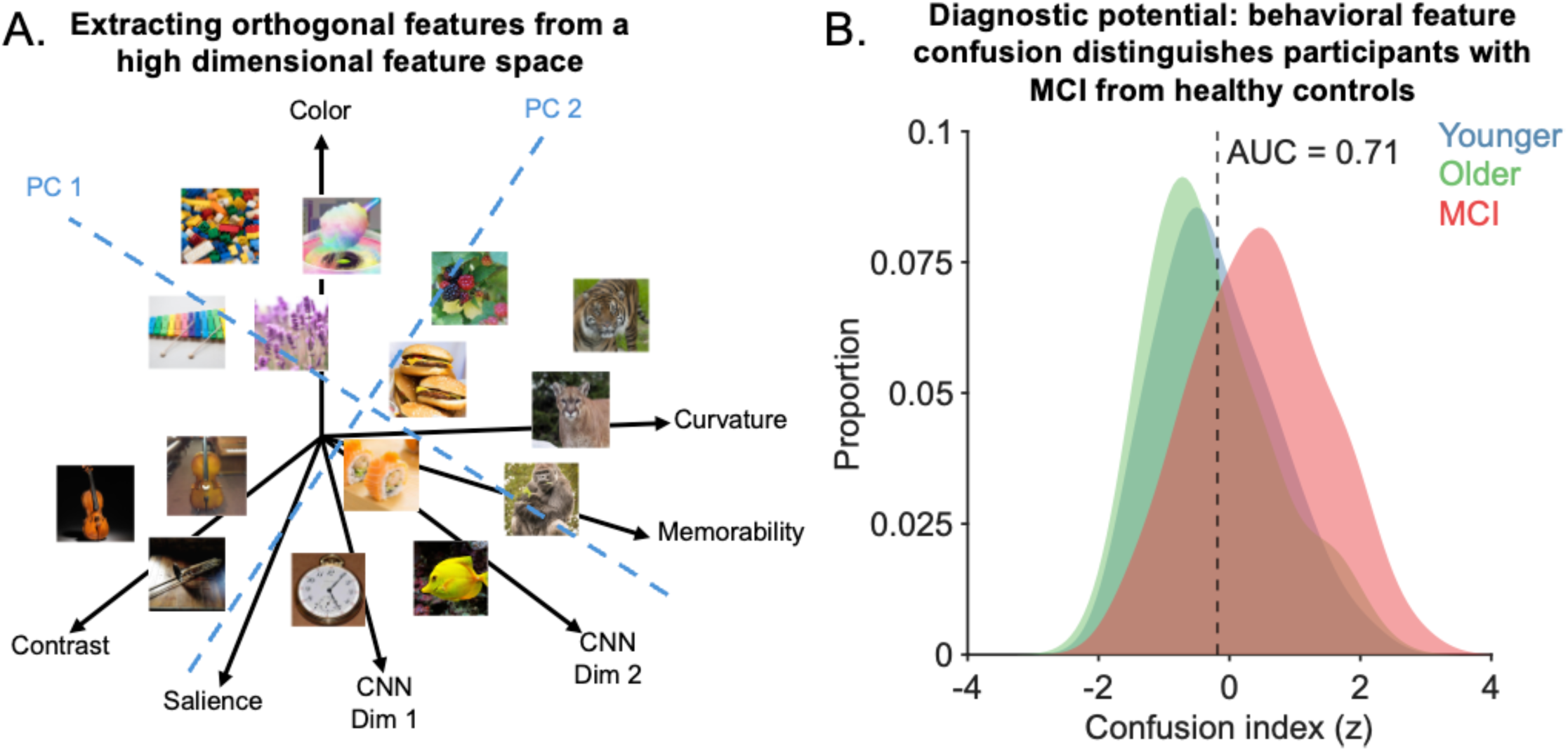
Behavioral feature confusion reflects cognitive health. (A) Schematic illustrating example dimensions in the richer high-dimensional image feature space. We extracted multiple properties of each image, including the visual features shown in Figure 1B, psychophysics-derived features such as salience and memorability^44,45,51^ (Supplemental Figure 3), and layer 5 activations of a deep convolutional neural network trained for image classification (VGG-16^50^). We then performed principal components analysis on this expanded image feature set to produce a set of 8 orthogonal feature dimensions with which to compute behavioral feature confusion. (B) Behavioral feature confusion distinguishes healthy controls from participants with MCI. We trained a logistic regression model to predict each participant’s image choices from orthogonalized features and pairwise interactions. As in Figure 3, we defined behavioral feature confusion index as the ratio of the root mean square weights of the interaction terms to the linear terms. Histograms show the resulting confusion index distributions for healthy controls and participants with MCI (n = 155 healthy vs. 83 MCI subjects, mean difference = -0.71, p = 6.7 x 10^-8^). Receiver operating characteristic (ROC) analysis on these indices distinguished participants with MCI from healthy controls with an area under the curve of 0.71 (n = 155 healthy vs. 83 MCI subjects, p = 1.9 x 10^-7^).

Participants with mild cognitive impairment showed stronger behavioral feature confusion than healthy controls (Figure 4B). Thus, the same phenomenon observed in neuronal populations and monkey behavior was also evident in a richer feature space incorporating perceptual and semantic properties of images.

Using this richer feature space, behavioral feature confusion distinguished individuals with mild cognitive impairment from healthy controls (area under the receiver operating characteristic curve comparing distributions for healthy controls and participants with MCI = 0.71). These results suggest that behavioral feature confusion is a signature of early cognitive impairment that generalizes across species and levels of description.

## Discussion

Early detection of Alzheimer’s disease has largely focused on identifying molecular pathology or measuring behavioral impairments after cognitive decline becomes apparent. Here we describe a complementary approach grounded in neuronal population representations. Across longitudinal measurements in a primate model of Alzheimer’s disease, we identified a change in population-level visual representations in which representations of distinct visual features become progressively less independent. This alteration had predictable consequences for behavior: preferences associated with one visual feature increasingly influenced choices associated with other, independent features. Using an analogous behavioral task in humans, we identified the same behavioral signature in individuals with mild cognitive impairment.

These findings establish a direct link between a measurable property of neuronal population activity and a behavioral signature associated with cognitive impairment. Importantly, this approach does not rely on overall task performance or memory failure. Instead, it focuses on structured relationships between visual information and behavior. Relationships among visual features revealed systematic changes in behavior that were not apparent in standard behavioral summaries or in the dominant sources of behavioral variation across individuals. Together, these results suggest that behavioral measurements grounded in neuronal population activity may provide a principled basis for detecting early functional changes associated with ADRD.

### Loss of independence in feature representations is reflected in behavior

Our results identify a change in population-level visual representations that emerges during disease progression and has predictable consequences for behavior. Importantly, disease progression did not eliminate visual feature information itself. Individual V4 neurons remained tuned to visual features throughout disease progression. Instead, the dominant change was a progressive loss of independence among feature representations. Visual features that were represented independently early in disease progression increasingly influenced one another at the population level, despite continued representation of the individual features themselves. In monkeys, neuronal feature confusion was accompanied by behavioral feature confusion. Humans with mild cognitive impairment showed the same behavioral signature. Thus, the present results demonstrate that neuronal population measurements in monkeys can guide the interpretation of structured behavioral changes in humans.

Several observations suggest that neuronal feature confusion does not arise primarily from degeneration of local feature encoding circuitry within V4. Tau pathology in V4 was sparse and was observed predominantly in processes rather than local neuronal cell bodies^10^. In addition, feature tuning at the level of individual neurons remained relatively stable throughout disease progression. This stability of visual feature tuning by individual units in V4 indicates that feedforward processes in the ventral stream likely remain intact early in disease progression. Together, these observations suggest that disease progression altered relationships among representations rather than eliminating local feature selectivity within V4.

One possible explanation is that neuronal feature confusion reflects disruption of feedback signals that normally help maintain separation among representations in visual cortex. Visual representations are shaped not only by feedforward sensory inputs, but also by signals related to attention, memory, task demands, behavioral context, and higher level visual processing^40,52–54^. Previous work from our laboratory has shown that the independence of feature representations can change transiently as a function of cognitive state, such that uncertainty about task demands increases interference between representations of unrelated visual features and produces corresponding behavioral confusion^1,2^. The present results raise the possibility that ADRD produces a progressive disruption of the same broader mechanisms that normally regulate or stabilize representational independence through disruption of feedback corticocortical circuits.

We focused on the visual system because decades of work in systems neuroscience have established detailed links between visual features, neuronal population activity, and visually guided behavior^36–41^. This foundation made it possible to identify a subtle alteration in neuronal population representations and predict corresponding changes in behavior. However, patients with ADRD often report forms of confusion in more cognitive domains, including mixing up details of distinct memories, confusing people or events, or calling family members by the wrong name^3–7^. Brain areas implicated in these functions, including entorhinal cortex, hippocampus, and frontal cortex, are affected early in disease progression in this monkey model and in ADRD in people^8,9,55^. An intriguing possibility is that disease-related alterations in neuronal population representations analogous to those observed here in visual cortex also emerge in these areas and contribute to these cognitive symptoms. If so, future work may make it possible to design sensitive behavioral tasks capable of detecting subtle and early behavioral signatures of these alterations in other domains of cognition.

Together, these findings demonstrate that neuronal feature confusion associated with disease progression is reflected in behavior in both monkeys and humans. This provides a basis for identifying early functional effects of disease through behavior, even when those effects are not captured by standard cognitive assessments.

### Implications for behavioral detection of early-stage disease

These findings suggest a complementary approach to detecting early-stage disease. Current diagnostic strategies rely largely on neuropsychological assessments and biological markers^15–24^. Biomarkers can provide sensitive indicators of pathology but are often invasive, expensive, and difficult to deploy broadly. Behavioral assessments typically focus on domains such as memory and executive function, where measurable deficits often emerge only after substantial disease progression^11,22,30^.

Simple visually guided behaviors offer a different type of measurement. These behaviors are straightforward to measure, can be precisely controlled, and depend on well-characterized relationships between visual input, neuronal population activity, and behavior^36–41^. Previous work has shown that image selection and exploration behavior are sensitive to cognitive impairment^1,2^, but commonly used measures such as novelty preference provide relatively coarse behavioral summaries^46–49,56^.

The approach presented here measures a different aspect of behavior. Rather than focusing on overall task performance or broad behavioral preferences, it quantifies relationships among the influences of multiple visual features on behavior. This makes it possible to detect consistent changes in how visual features jointly influence choice, even when overall patterns of image selection remain largely preserved. In both monkeys and humans, these changes emerged early and were detectable using a simple task that required minimal training.

Because this approach is grounded in quantitative relationships between visual features, neuronal population activity, and behavior, it provides access to aspects of behavior that are not captured by standard assessments. The use of visually guided tasks also supports scalability, as these measurements can be implemented in brief online formats and repeated longitudinally. Repeated measurements within individuals may make it possible to derive continuous measures sensitive to subtle changes over time. These findings therefore suggest that behavioral assays grounded in neuronal population representations may provide a scalable and non-invasive route for detecting early functional effects of disease.

### A primate model of disease provides a basis for interpreting behavior in humans

ADRD research has focused largely on two complementary goals: identifying the biological mechanisms associated with disease and characterizing how cognition changes in human patients. Molecular and cellular studies have identified many of the pathological processes associated with disease progression^15–21^, while human studies have described alterations in memory, executive function, and behavior^22–24^. However, connecting these levels of analysis remains difficult. Biological mechanisms do not influence behavior directly. Their effects must ultimately be expressed through changes in neuronal population representations that influence behavior.

The present results suggest that neuronal population representations provide a critical intermediate level for linking biological changes to cognition. In this study, disease progression altered relationships among visual feature representations in neuronal populations and produced corresponding changes in visually guided behavior. Because these relationships could be measured quantitatively, they made it possible to generate specific behavioral predictions from changes in neuronal population representations and to test those predictions in humans.

A primate model of disease provides experimental access to this level of description. Because primates exhibit sophisticated visually guided behaviors supported by well-characterized visual systems, primate models uniquely allow changes in neuronal population representations to be related directly to behavior^57–59^. In the present study, neuronal population activity and behavior were measured longitudinally in the same individuals before and during disease progression. This made it possible to identify changes in neuronal population representations and relate those changes directly to measurable changes in behavior within individuals over time. Such relationships are difficult to establish using molecular measurements or behavioral measurements alone.

This framework also provides a basis for interpreting behavior in humans in biological terms. Although neuronal population activity cannot readily be measured longitudinally at comparable resolution in human patients, behavior can be analyzed using the same relationships that link neuronal population representations to behavior in the model system. In this way, measurements in the primate model can guide the interpretation of subtle behavioral changes in humans and provide a framework for connecting biological mechanisms to cognition through neuronal population representations.

### Next steps

These findings demonstrate that behavior can reveal subtle disease-related changes in neuronal population activity before impairments are detectable using standard cognitive assessments. Because these behavioral measurements are grounded in relationships between neuronal population representations and behavior, they provide more than a general measure of impairment: they provide information about how disease alters the neuronal computations that generate behavior.

This creates an opportunity to detect and track early functional changes associated with ADRD using behavioral measurements that are rapid, scalable, and non-invasive. In the present work, even a single session of a simple online task distinguished individuals with mild cognitive impairment from healthy controls. Integrating behavioral measurements of this kind with molecular biomarkers, neuroimaging, and genetic risk factors could improve early detection of disease-related brain changes and help identify patients before substantial cognitive decline emerges.

These approaches may also prove useful for evaluating treatments. Current outcome measures often focus on broad behavioral impairment or molecular pathology, making it difficult to determine whether interventions alter the neuronal computations that generate behavior. Behavioral measurements grounded in neuronal population activity create an opportunity to assess whether treatments modify the functional changes linking pathology to behavior, potentially enabling more sensitive tracking of disease progression and therapeutic efficacy over time.

Together, these results identify a specific alteration in neuronal population representations that links disease progression to behavior across species. They demonstrate that disease-related changes in brain activity can leave measurable and biologically interpretable signatures in behavior, creating new opportunities to detect, track, and potentially intervene on early functional changes associated with Alzheimer’s disease.

## Methods

### Animals

Two adult male rhesus macaques (10 to 12 years old) had bilateral stereotaxic injections of an adeno-associated virus expressing a double tau mutation (AAV-P301L/S320F, 1.176 × 10^13 genomic copies/mL) into entorhinal cortex^8,9^. To increase the extent of AAV expression, two injections were made per hemisphere. Injections were performed using the StealthStation surgical navigation system (Medtronic, Minneapolis, MN) using presurgical MRIs as guidance. During the same surgery, and following the injections, two 6×8 microelectrode arrays (Blackrock Microsystems) were implanted subdurally in the left hemisphere of each animal in visual area V4. The two arrays were connected to each of two percutaneous connectors, which allowed electrophysiological recordings. The distance between adjacent electrodes was 400 μm, and each electrode was 1 mm long. Biofluid samples were collected from both animals immediately prior to surgery, approximately every 30 days following the AAV injections, as well as immediately prior to euthanasia. All experiments were conducted with approval by the Institutional Animal Care and Use Committee (IACUC) at the University of California–Davis.

### AAV preparation

The adeno-associated virus used in this study has been described in detail previously^8,9,60^. A capsid 1 adenovirus was packaged with one copy of human 0N/4R tau-containing two mutations (P301L/S320F) that render it more aggregation-prone than wild-type tau. The expression of this tau insertion is under the control of the hybrid cytomegalovirus enhancer/chicken β-actin (CMV/CBA) promoter, a CBA intron (first intron of chicken β-actin gene plus the splice acceptor of the rabbit β-globin gene), woodchuck hepatitis virus post-transcriptional regulatory element (WPRE), and bovine polyA. Virus packaging and purification were performed at the Penn Vector Core (University of Pennsylvania), following Good Laboratory Practices, including endotoxin testing (<5 endotoxin units per mL), purity testing (sodium dodecyl sulfate-polyacrylamide gel electrophoresis), and titration (three rounds of digital polymerase chain reaction). After preparation, aliquots of AAVs were stored at −80°C until surgery.

### Structural MRI and surgical planning

We imaged the monkeys’ brains using a 3T MRI scanner (Skyra, Siemens Healthcare, Germany) with an 8-channel receiver coil optimized for structural monkey brain scanning (Rapid MRI, Columbus, OH). T_1_-weighted images were acquired using the magnetization-prepared rapid acquisition with gradient echo (MPRAGE) pulse sequence with the following parameters: field of view = 154×154 mm; 480 sagittal slices; TR/TE = 2500/3.65 ms; flip angle = 7°; TI = 1100 ms; voxel size 0.6×0.6×0.6 mm, interpolated to a resolution of 0.3×0.3×0.3 mm. Anatomical images were used to identify the location of the entorhinal cortex and, in conjunction with fiducial markers, determine appropriate injection sites.

### Animal observation and health monitoring

We tracked both animals’ weights weekly and observed slight increases over the year of post-surgical testing. Veterinary, animal care, and laboratory staff performed regular physical exams, monitored food intake, and conducted informal assessments of cage-side behavior. Both animals maintained stable relationships with their pair-housed partners and frequently engaged in social interactions and grooming. Neither animal showed major behavioral changes that required intervention. Throughout the testing period, both animals continued to voluntarily enter the primate chair and perform the task for juice and food rewards, although the number of completed trials per session gradually declined in both cases.

### Behavioral training and testing in monkeys

We seated each monkey in a primate chair facing a computer monitor and provided access to a rigid straw for liquid rewards. During early training, we positioned padded spacers beside the head to encourage a forward-facing posture, but we removed them before recording sessions. We tracked eye position (using Eyelink infrared eye tracker) and only advanced trials when the monkey faced forward and placed its mouth on the reward straw. Before surgery, each monkey trained for 3–6 months on a set of visually guided tasks. In the home cage, they had ad libitum water; during experiments, we delivered juice or food rewards.

In the image selection task, each trial began when the monkey fixated a central target for 250 ms. We then presented two images in opposite hemifields. During recordings, one image was placed in the joint receptive fields of the V4 neurons. The monkey maintained fixation for another 150 ms before the fixation point disappeared, signaling the start of free viewing. To introduce new images, we presented the same novel image both inside and outside the receptive field and allowed 1500 ms of free viewing. Once an image became familiar (i.e. presented more than once), it was either paired with a truly novel image or with itself, and the free-viewing period lasted 1000ms in both cases. We drew images pseudo-randomly from the THINGS image set^44^ and manipulated familiarity so that each stimulus was viewed between one and hundreds of times. Over the course of approximately 13 months, Monkey C viewed 3,358 unique images and Monkey M viewed 2,415. We defined the selected image as the one the monkey first fixated during the free viewing period. Sessions were included for behavioral analyses if a minimum number of trials were successfully completed (20 trials for both Monkey C and Monkey M). This resulted in 63 sessions with an average of 38 trials per session for Monkey C, and 47 sessions with an average of 34 trials per session for Monkey M for behavioral analyses.

### Description of human subjects

We recruited 239 adult participants through Prolific (prolific.co), all of whom performed an adapted version of the image selection task (described below). All participants reported being at least 18 years old, had normal or corrected to normal vision, and spoke English fluently. We classified young adults as healthy individuals 18-30 years old, older adults as 60-84 years old, and MCI adults as adults of any age who self-reported as being diagnosed with MCI (actual age range 19-75). Each subject completed a survey containing questions about demographics and general lifestyle prior to beginning the experiment. There were 79 young adults (44 male, 26 female, 9 other), 76 old adults (26 male, 46 female, 4 other), and 84 adults with MCI (21 male, 50 female, 13 other). Across all groups, subjects were predominantly white or black/African American (51% white, 43% black/African American, 4% Asian, 2% other) and lived across 19 countries around the world. Additional survey questions assessed lifestyle variables including drinking, smoking, exercise, social engagement, creative activities, video game use, television viewing, stimulant use, and reproductive health measures, as well as diagnosis of attention-deficit/hyperactivity disorder and family history of Alzheimer’s disease. Including the survey and completing 900 trials of the task, subjects typically completed the experiment in approximately one hour. All procedures were approved and granted a type 4 exemption by the Institutional Review Board of the University of Chicago.

### Behavioral task for humans

We adapted the monkey image selection task for online testing with human participants. Each trial began when the participant clicked a central fixation cross. Two images then appeared side by side for 150 ms, after which they disappeared and a solid mask covered both locations. Participants clicked on the image they preferred to see again. The chosen image was then revealed for 1000 ms.

As in the monkey version, some trials included two identical images, whereas others paired one familiar image with a novel image. Images were viewed between one and 54 times. Across the task, each participant viewed 900 images in total.

### Neuronal recordings

We recorded neuronal activity from the electrode arrays during daily experimental sessions over one year in each monkey. Using our recording methods, it is nearly impossible to tell whether we recorded from the same single- or multi-unit clusters on subsequent days. To be conservative, all analyses were performed independently within single recording sessions and then compared across time when relevant.

Two 64-channel multielectrode arrays were chronically implanted in visual area V4. The threshold for each channel was set at three times the standard deviation and the threshold crossings were used as multiunit activity for that channel. The stimuli were positioned over the joint receptive fields of the V4 units from both arrays, which were always in the left hemisphere. Sessions were included for neural analyses if the total number of completed trials was at least 20.

Neuronal responses were typically quantified as spike count responses from 60-210 ms after stimulus onset. This is the stimulus presentation period while the monkey fixated the central point, using a window shifted by 60 ms to account for V4 response latency^61^.

We use the term “unit” to refer to multiunit activity recorded on one electrode. Analyses comparing single units and multiunits in our previous work^62^ as well as from another laboratory^63^ did not find systematic differences between single neurons and multiunits for the types of neuronal analyses presented here. V4 units were included for analysis if the stimulus-evoked response was sufficiently larger than baseline activity (75% greater for Monkey C, 100% greater for Monkey M), measured 100 ms before stimulus onset. Sessions were included for analysis if a minimum number of responsive units were recorded (10 units for both Monkey C and Monkey M) as well as if a minimum number of successful trials were completed (20 trials for Monkey C, 15 trials for Monkey M). These inclusion criteria resulted in 84 sessions from Monkey C with an average of 63 trials and 96 V4 units per session, and 64 sessions from Monkey M with an average of 43 trials and 41 V4 units per session for neuronal analyses.

### Data analysis and statistical comparisons

Changes in neuronal and behavioral measures over time were pooled across the two monkeys and assessed as the Pearson correlation between each measure and day against the null hypothesis of a correlation of zero. The same approach was used to assess relationships between neuronal population encoding axes of visual image features (e.g., contrast, curvature, Figure 1C). To assess differences in the distribution of behavioral measurements between groups of human subjects, we used two-sided two-sample t-tests. The quantification of discriminability of health controls and MCI subjects was assessed as the area under the receiver operating characteristic curve.

### Extraction of image-computable visual features

Before extracting image-computable visual features, we preprocessed the images to standardize image size and facilitate feature extraction. We rescaled each image to 800 × 800 pixels and converted images to grayscale before extracting grayscale-based features using built-in MATLAB functions (MathWorks 2026a).

We extracted a broad set of image-computable visual features describing luminance statistics, spatial structure, texture, shape, color, and local image features (Supplementary Fig. 1). These included contrast (pixel standard deviation); orientation (average vector of gradient directions); curvature (standard deviation of the Laplacian transform); sharpness (variance of the Laplacian transform); skewness (pixel intensity asymmetry); kurtosis (pixel intensity tailedness); Fourier energy (average fast Fourier transform magnitude); red, green, and blue color statistics (average normalized pixel intensity per channel); texture contrast, correlation, energy, and homogeneity (gray-level co-occurrence matrix statistics); symmetry (difference between the image and its mirrored version); edge density (fraction of edge pixels detected using a Sobel operator); aspect ratio and bounding box size (calculated from the largest object in the binarized image); corner count (Harris corner detector); histogram of oriented gradients (HOG) length; ORB keypoint count (Oriented FAST and Rotated BRIEF); and SURF blob count (Speeded-Up Robust Features).

### Extraction of psychophysics-derived image features

To complement the image-computable visual features described above, we extracted several higher-level features from the THINGS images that were derived from human psychophysics experiments and computational models. We included these features to improve discriminability between healthy participants and individuals with MCI.

We extracted salience for each image using a deep learning convolution neural network^64^ trained to predict image salience from natural image statistics and human gaze patterns. The network generated salience values for each pixel, which we averaged and normalized to obtain a single salience value per image.

The remaining features were derived from published human psychophysics datasets. Memorability scores came from more than one million trials in which participants judged whether images were novel or previously viewed^51^. Arousal scores came from an experiment where participants were asked to rate images from 1 (very calming) to 7, (very arousing)^65^. Perceptual similarity scores were derived from an odd-one-out task in which participants were asked to select which of three images was most dissimilar from the others^45^. We applied multidimensional scaling to the resulting dissimilarity matrix and used the dominant dimension of the embedding to assign each image a scalar similarity score reflecting how similar it was to other THINGS images.

### Orthogonalization and dimensionality reduction of image-computable visual features

For neuronal feature confusion analyses in monkeys, we applied principal components analysis (PCA) to the matrix of image-computable visual features within each session. This transformed the original correlated feature set into orthogonal feature dimensions and reduced dimensionality (10 PCs for Monkey C, 7 PCs for Monkey M). We then z-scored each feature dimension to normalize differences in variance across dimensions.

For behavioral feature confusion analyses in monkeys and humans, we applied PCA to the matrix of differences in image-computable visual features between the right and left images within each session. As above, PCA transformed correlated visual features into orthogonal feature dimensions while reducing dimensionality (10 PCs for Monkey C, 5 PCs for Monkey M, 5 PCs for humans). We z-scored each feature dimension before subsequent analyses.

### Estimation of neuronal feature confusion in monkeys

For each recording session and each feature dimension, we used linear regression to predict feature values from the vector of neuronal responses. We performed leave-one-out cross-validation across trials within each session by training the model on all trials except one and predicting the feature value for the held-out trial. Repeating this procedure across trials produced a predicted value for each trial and feature dimension. We quantified decoding accuracy as the correlation coefficient (r) between predicted and actual feature values across trials within each session.

To quantify neuronal feature confusion, we performed a cross-decoding analysis in which we used the linear model fit to one feature dimension to predict the values of other feature dimensions. For each pair of feature dimensions, we normalized cross-decoding performance by the average self-decoding performance of the two feature dimensions. We included only feature dimensions with self-decoding accuracy above threshold (r = 0.1 for Monkey C, r = 0.05 for Monkey M). For each session, we calculated average neuronal feature confusion across all feature pairs and then z-scored the resulting values across sessions. Higher neuronal feature confusion indicates greater overlap in neuronal population representations across distinct feature dimensions.

### Image-computable visual feature correlations with image selection

For each trial, we calculated the relationship between image choice and the difference in each image-computable visual feature between the two presented images (left − right; Figure 2B). For monkeys, image choice was defined by the first saccade during free viewing. For humans, image choice was defined by mouse click. We visualized these feature–behavior relationships for individual features over time (Figure 2C, monkeys) and across all features (Figure 2D, monkeys across time; Figure 3B, humans across subjects). Positive correlations (yellow in Figure 2D and Figure 3B) indicate that subjects tended to select images with larger values of a given feature, whereas negative correlations (blue in Figure 2D and Figure 3B) indicate that subjects tended to select images with smaller values of that feature.

### Estimation of behavioral feature confusion in monkeys

Behavioral feature confusion quantifies the extent to which behavior driven by one visual feature predicts behavior driven by other, independent features. For each session, we fit a separate logistic regression model predicting image choice from each feature dimension and defined that dimension’s self-prediction as the correlation between predicted and actual choice. For each pair of dimensions, we defined cross-prediction as the magnitude of the correlation between their predicted choices . We normalized each pair’s cross-prediction by the mean of the two dimension’s self-predictions. We restricted this to pairs in which both dimensions individually predicted choice above 0.05 and excluded sessions with fewer than 10 pairs. Behavioral feature confusion for a session was the mean normalized value across included pairs, z-scored across sessions.

### Estimation of behavioral feature confusion in humans

In the human experiments, we had enough trials to define behavioral feature confusion as the relative importance of individual features and their interactions in predicting each participant’s decision. To compute behavioral feature confusion for image features (Figure 3), we characterized each image using the same orthogonalized image features extracted previously for the analogous monkey analysis. For confusion based on the richer image set (Figure 4), we additionally included a broad set of properties, including psychophysics-derived measures (memorability, salience, arousal, perceptual similarity) and activations from a deep convolutional neural network. In both cases, many of these features are correlated in natural images, so we used PCA to identify orthogonal feature dimensions. This transformation produced feature dimensions that were independent in image space, allowing interactions among features to be distinguished from correlations in the images themselves.

For each trial, we computed the difference between the projections of the left and right images along each principal component. We fit a logistic regression model for each participant that predicted image choice from those differences in feature dimensions (linear terms) and their interactions. Because the principal components were orthogonal in image space, pairwise interaction terms captured dependencies in how feature dimensions influenced behavior. We defined behavioral feature confusion as the logarithm of the ratio of the root mean square of the weights of interaction and linear terms. Larger values indicate stronger interactions among feature dimensions.

## Acknowledgements

We thank Philip Sulewski for discussions and comments on an earlier version of the manuscript and Kursti Ropp Eaves for technical assistance.

## Author contributions

DAR, MRC, and JHM conceptualized the monkey experiment. DEGS, KSA, and MRC conceptualized the human experiment. DAR performed the monkey experiments. DEGS performed the human experiments. DEGS, DAR, RS, and MRC performed data analysis. JHM and MRC supervised and funded the work. All authors wrote the paper and provided comments and discussion about the manuscript.

## Materials & Correspondence

Correspondence and material requests should be addressed to MRC.

