## Supplemental Information for "Progressive loss of independence in neuronal representations predicts cognitive decline"

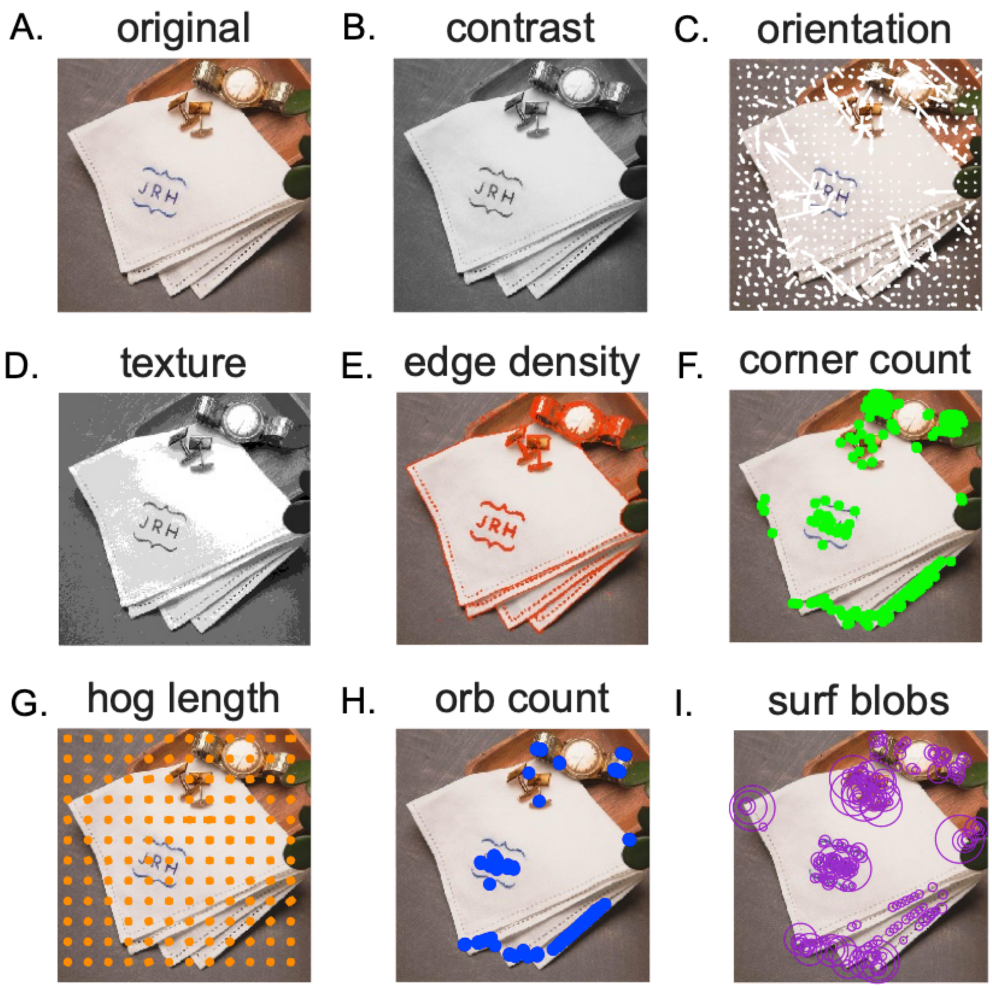


***Supplemental Figure 1.*** *Visualizations of example image-computed visual features. Each image was preprocessed and underwent a series of transformations to extract visual feature values. While a single value was computed for each feature from each image, some features (contrast, orientation, texture-related features, edge density, corner count, histogram of oriented gradients, oriented FAST and rotated BRIEF keypoints, SURF keypoints) had an intermediate step that lent itself to visualization (see Methods for details on each feature). A) Original image. B) Grayscale image from which contrast is computed. C) White arrows denote local gradient direction at each sampled point and are aligned to the direction of steepest intensity change from which global orientation is calculated. D) Texture-related features (texture contrast, texture correlation, texture energy, texture homogeneity) are derived from the gray-level co-occurrence matrix (overlayed on the image) which denotes varying intensity between neighboring pixels. E) Red denotes pixels classified as edges by a Sobel edge detector. F) Green denotes Harris corner keypoints. G) Orange denotes oriented gradient bars derived from local shape and texture structure. H) Blue marks ORB keypoints where binary features cluster around local structures. I) Purple denotes SURF keypoints, circle size denotes the spatial scale at which each blob was detected.*

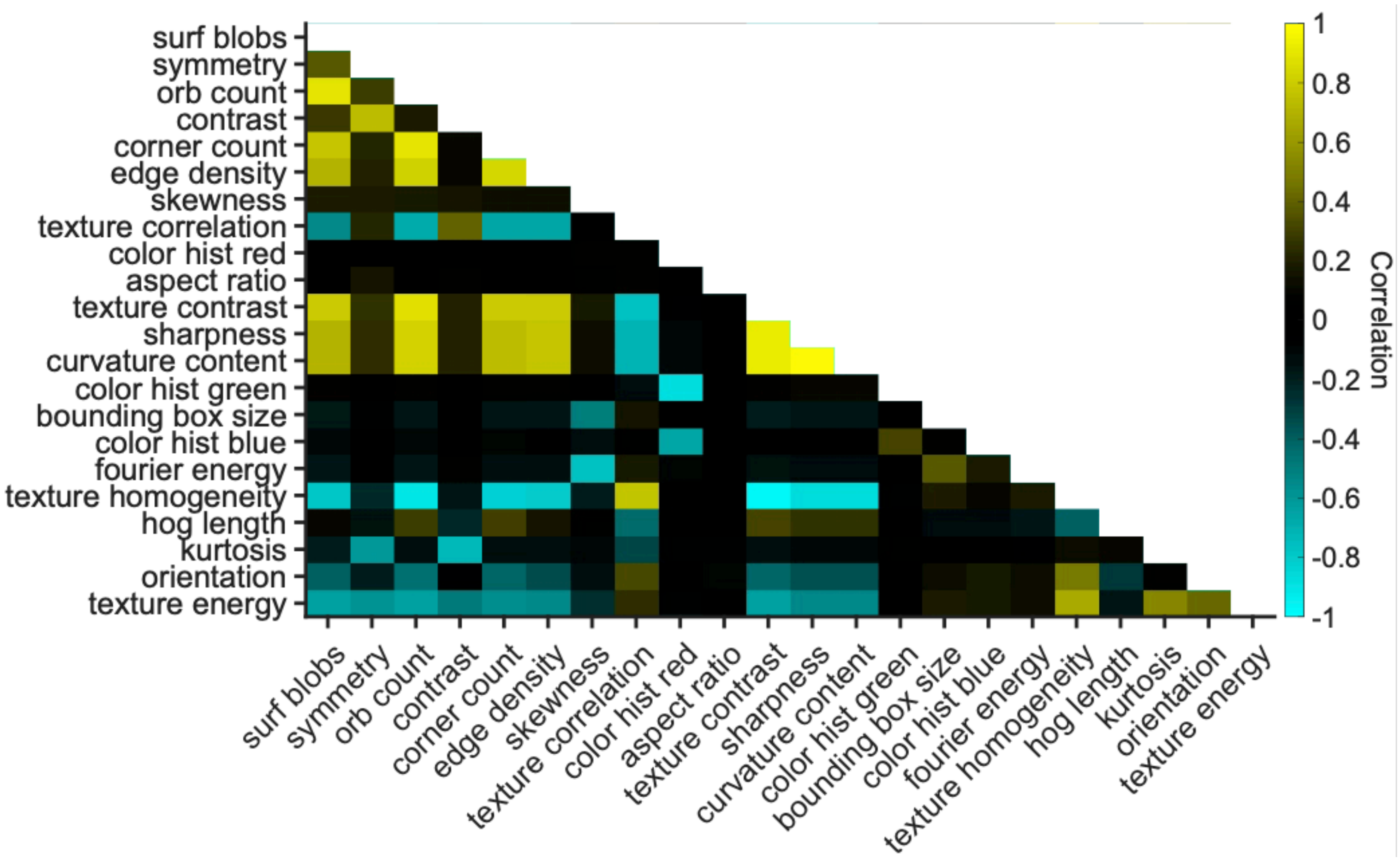


***__Supplemental Figure 2.__*** *Correlations between values of image-computable visual features across the THINGS image set. Yellow colors indicate positive correlations, and cyan colors indicate negative correlations. Some pairs of image features are strongly correlated while others are largely independent.*

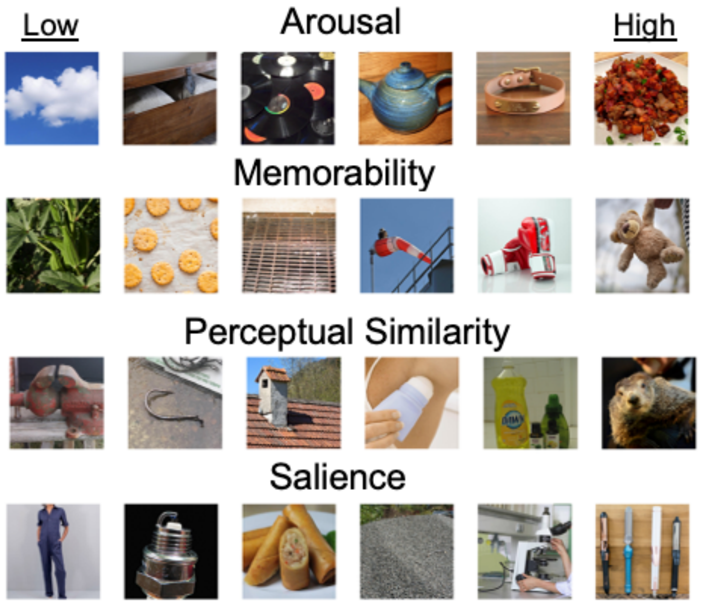


***Supplemental Figure 3.*** *Example psychophysics-derived features. Example images spanning the range of the four features: arousal, memorability, perceptual similarity, and salience (see Methods for feature details). Images span an evenly spaced gradient of the feature space from lowest (left) to highest (right) feature values.*
